# Leveraging genetic interaction for adverse drug-drug interaction prediction

**DOI:** 10.1101/455006

**Authors:** Sheng Qian, Siqi Liang, Haiyuan Yu

## Abstract

In light of increased co-prescription of multiple drugs, the ability to discern and predict drug-drug interactions (DDI) has become crucial to guarantee the safety of patients undergoing treatment with multiple drugs. However, information on DDI profiles is incomplete and the experimental determination of DDIs is labor-intensive and time-consuming. Although previous studies have explored various feature spaces for *in silico* screening of interacting drug pairs, no method currently provides reliable predictions outside of their training sets. Here we demonstrate for the first time targets of adversely interacting drug pairs are significantly more likely to have synergistic genetic interactions than non-interacting drug pairs. Leveraging genetic interaction features and a novel training scheme, we construct a gradient boosting-based classifier that achieves robust DDI prediction even for drugs whose interaction profiles are completely unseen during training. We demonstrate that in addition to classification power—including the prediction of 432 novel DDIs—our genetic interaction approach offers interpretability by providing plausible mechanistic insights into the mode of action of DDIs.

## INTRODUCTION

Drug-drug interactions (DDIs) refer to the unexpected pharmacologic or clinical responses due to the co-administration of two or more drugs ^1^. With the simultaneous use of multiple drugs becoming increasingly prevalent, DDIs have emerged as a severe patient safety concern over recent years ^2^. According to The Center for Disease Control and Prevention (CDC), the percentage of Americans taking three or more prescription drugs in the past 30 days increased from 11.8% in 1988-1994 to 21.5% in 2011-2014, and the occurrence of polypharmacy, defined as the concurrent use of five or more drugs, increased from 4.0% to 10.9% within the same time period ^3,4^. Polypharmacy is especially common among elderly people, affecting 42.2% of Americans aged 65 years and older, exposing them to a higher risk of adverse DDIs. Indeed, DDIs were estimated to be responsible for 4.8% of hospitalization in the elderly, a 8.4-fold increase compared to the general population ^5^. Overall, DDIs contribute to up to 30% of all adverse drug events (ADEs) ^6^ and account for about 74,000 emergency room visits and 195,000 hospitalizations each year in the United States alone ^3^. Therefore, it has become a medical imperative to identify and predict interacting drug pairs that lead to adverse effects.

In order to facilitate identification of interacting drug pairs, a number of *in vitro* and *in vivo* methods have been developed. For example, drug pharmacokinetic parameters and drug metabolism information collected from *in vitro* pharmacology experiments and *in vivo* clinical trials can be used to predict interacting drug pairs ^7,8^. However, these methods are labor-intensive and time-consuming, and are thus not scalable to all unannotated drug pairs ^9^. In the past decade, machine learning-based *in silico* approaches have become a new direction for predicting DDIs by leveraging the large amount of biological and phenotypic data of drugs available. The advantage of machine learning-based approaches lies in their ability to perform large-scale DDI prediction in a short time frame. So far, various features have been explored for building DDI prediction models, including similarity-based features and network-based features, among others. Similarity-based features characterize the similarity of the two drugs at question in terms of chemical structure, side effect profile, indication, target sequence, target docking, ATC group, etc. ^10–24^. Network-based features exploit the topological properties of the drug-drug interaction network or the protein-protein interaction network, which relates to DDIs through drug-target associations ^16,25–27^. While these methods have yielded important information about DDIs, few methods to date have been able to provide insight into the molecular mechanisms of drug-drug interactions.

To this end, in this study, we employ the genetic interaction between genes that encode the targets of two drugs as a novel feature for predicting interacting drug pairs that cause adverse drug reactions. We show that targets of adversely interacting drugs tend to have more synergistic genetic interactions than targets of non-interacting drugs. Exploiting this finding, we apply a machine learning framework (**Supplementary Fig. 1**) and build a gradient boosting-based classifier for adverse DDI prediction by integrating genetic interaction and three widely used features – indication similarity, side effect similarity and target similarity. We show that our model provides accurate DDI prediction even for pairs of drugs whose interaction profiles are completely unseen during training. Furthermore, we find that excluding the genetic interaction features significantly decreases the performance of our model. Through genetic interactions, our method provides insight into the mode of action of drugs that lead to adverse combinatory effects.

## RESULTS

### Genetic interaction profiles provide complementary information for distinguishing interacting and non-interacting drugs

In order to explore the separating power of various features to distinguish adversely interacting drug pairs from non-interacting drug pairs, we constructed a high-confidence set of adversely interacting drug pairs from all DDIs labeled “the risk or severity of adverse effects can be increased” in DrugBank ^28^ (**Supplementary Table 1**). This resulted in a set of 117,045 adversely interacting drug pairs involving 2,261 drugs. 2,195,023 non-interacting drug pairs were generated by taking all other combinations of these drugs before removing any drug pair that has been reported in DrugBank, TWOSIDES ^29^ or a complete dataset of DDIs compiled from a variety of sources ^30^. Furthermore, we required that all features, including indication similarity, side effect similarity, target sequence similarity and genetic interaction, should be available for each drug pair. After this filtering step, 1,113 adversely interacting drug pairs and 11,313 non-interacting drug pairs involving 262 drugs remained.

Interacting and non-interacting drug pairs exhibit different distributions in terms of the four groups of properties that we investigated. Indications and side effects of drugs were mapped to four levels of the MedDRA hierarchy ^31^ (**Fig. 1a**). At every level, adversely interacting drugs are associated with significantly more similar side effects as well as indications than non-interacting drugs (**Fig. 1b-c, Supplementary Fig. 2a-b**). On another front, target similarity was calculated by aligning the sequences of the protein targets with the Smith-Waterman algorithm ^32^. Since a drug may have multiple protein targets, aggregation was performed by taking the minimum, mean, median or maximum alignment score for each drug pair (**Fig. 1d**). As expected, the maximum, mean and median target similarity between targets of adversely interacting drug pairs are significantly higher than those of non-interacting drug pairs (**Fig. 1e**). Interestingly, interacting drug pairs manifest a significantly lower minimum target similarity than non-interacting drug pairs (**Fig. 1e**). This could be due to the fact that interacting drugs possess a higher number of protein targets combined, thereby having a higher change of targeting vastly different targets (**Supplementary Fig. 3a**). These results establish indication similarity, side effect similarity and target similarity as informative predictors of adverse DDIs.

**Figure 1.**
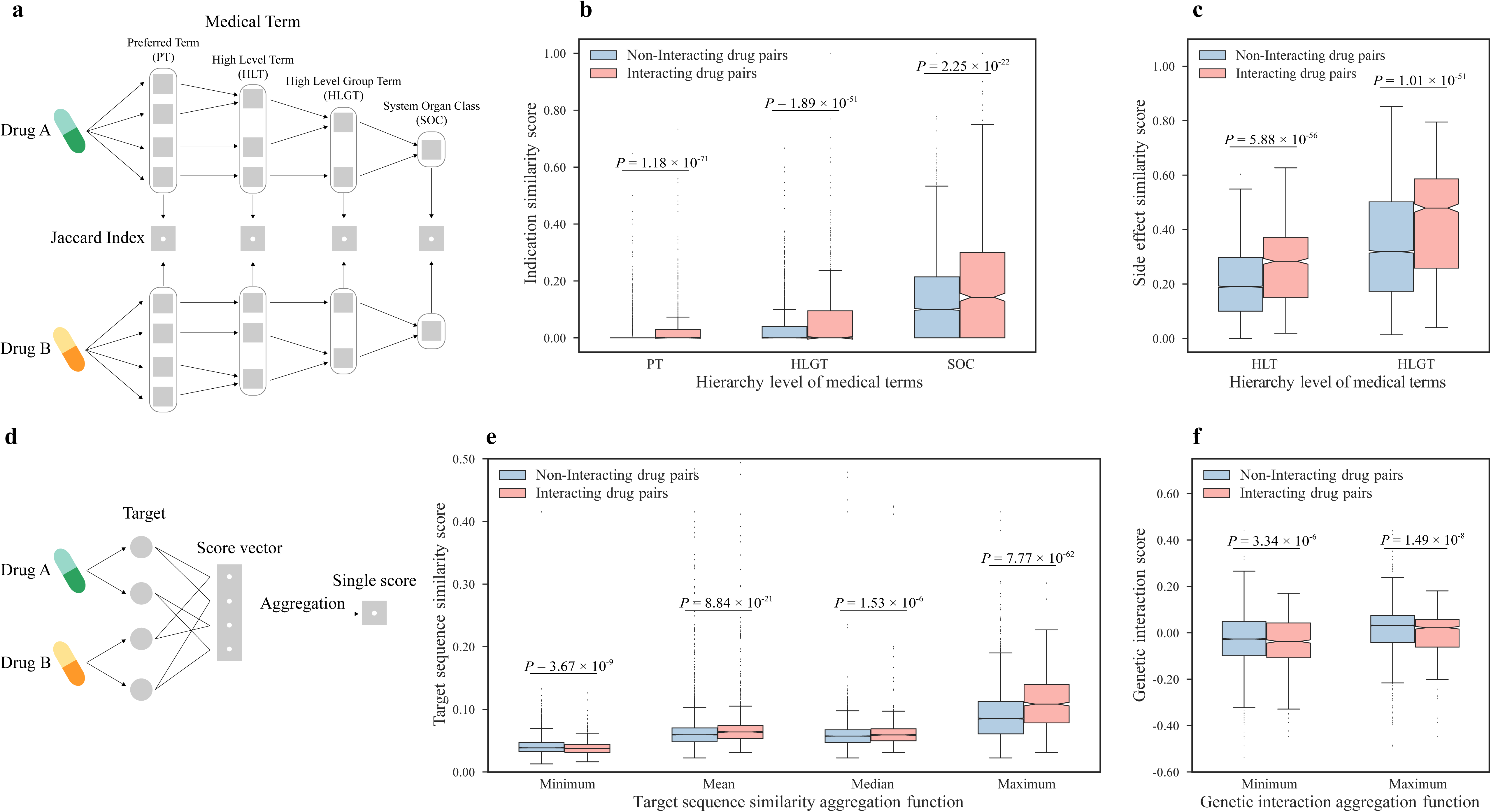
Adversely interacting drug pairs and non-interacting drug pairs significantly differ with regard to the 11 features selected. (a) Schematics of calculating indication similarity and side effect similarity features. (b) Indication similarity score of hierarchy level PT, HLGT and SOC between two drugs. (c) Side effect similarity score of hierarchy level HLT and HLGT between two drugs. (d) Schematics of calculating target sequence similarity and genetic interaction features. (e) Minimum, mean, median and maximum target sequence similarity score between targets of two drugs. (f) Minimum and maximum genetic interaction score between targets of two drugs. (Statistical significance determined by two-sided Mann-Whitney U test)

Genetic interaction refers to deviation from the expected phenotype when two genes are simultaneously mutated ^33^. The combined phenotype can be less severe than expected, as in the case of buffering interactions, or it can be more extreme than expected, as in the case of synergistic interactions ^34^. Since binding of drugs modulates the function of their targets, the genetic interaction between protein targets of two drugs might be associated with their joint effects. On this account, we investigated whether targets of adversely interacting drugs and targets of non-interacting drugs display divergent genetic interaction profiles. For each pair of drugs, we mapped their protein targets to the corresponding yeast homologs and obtained genetic interaction scores between the yeast genes from a global yeast genetic interaction network ^35^. When the minimum, mean, median or maximum genetic interaction score was taken for targets of each drug pair, adversely interacting drugs showed significantly lower scores than non-interacting drugs irrespective of the aggregation function applied (**Fig. 1f, Supplementary Fig. 2c**). This trend can be recapitulated using a recently published human genetic interaction dataset (**Supplementary Fig. 3b**). Furthermore, genetic interaction provides complementary information that is not captured by target similarity, indication similarity, or side effect similarity, as seen from their poor correlation (**Supplementary Fig. 4**). Therefore, genetic interaction profiles of drug targets provide new information as a predictor of adverse DDIs.

### Building a machine learning model for predicting adverse DDIs

To divide drug pairs into a training set and a test set for building a machine learning model, most previous studies randomly split their data with a specified ratio ^10,16,17,19,22,23,36,37^, without considering the fact that drugs appearing in both sets may carry extra information about their interaction propensity. Considering the scenario of predicting interactions of drugs without prior information about their interaction profiles, this splitting scheme becomes inappropriate. To address this problem, we draw on a method that partitions drug pairs based on drugs ^14,20,21^. All drugs in our constructed dataset were randomly split into “training drugs” and “test drugs” with a ratio of 2:1. The training set consists of all drug pairs where both drugs are “training drugs” and the test set comprises all drug pairs where both drugs are “test drugs” (**Fig. 2a**). As a result, 475 interacting drug pairs and 4,802 non-interacting drug pairs involving 175 drugs went into the training set; 131 interacting drug pairs and 1,322 non-interacting drug pairs involving 87 drugs went into the test set.

**Figure 2.**
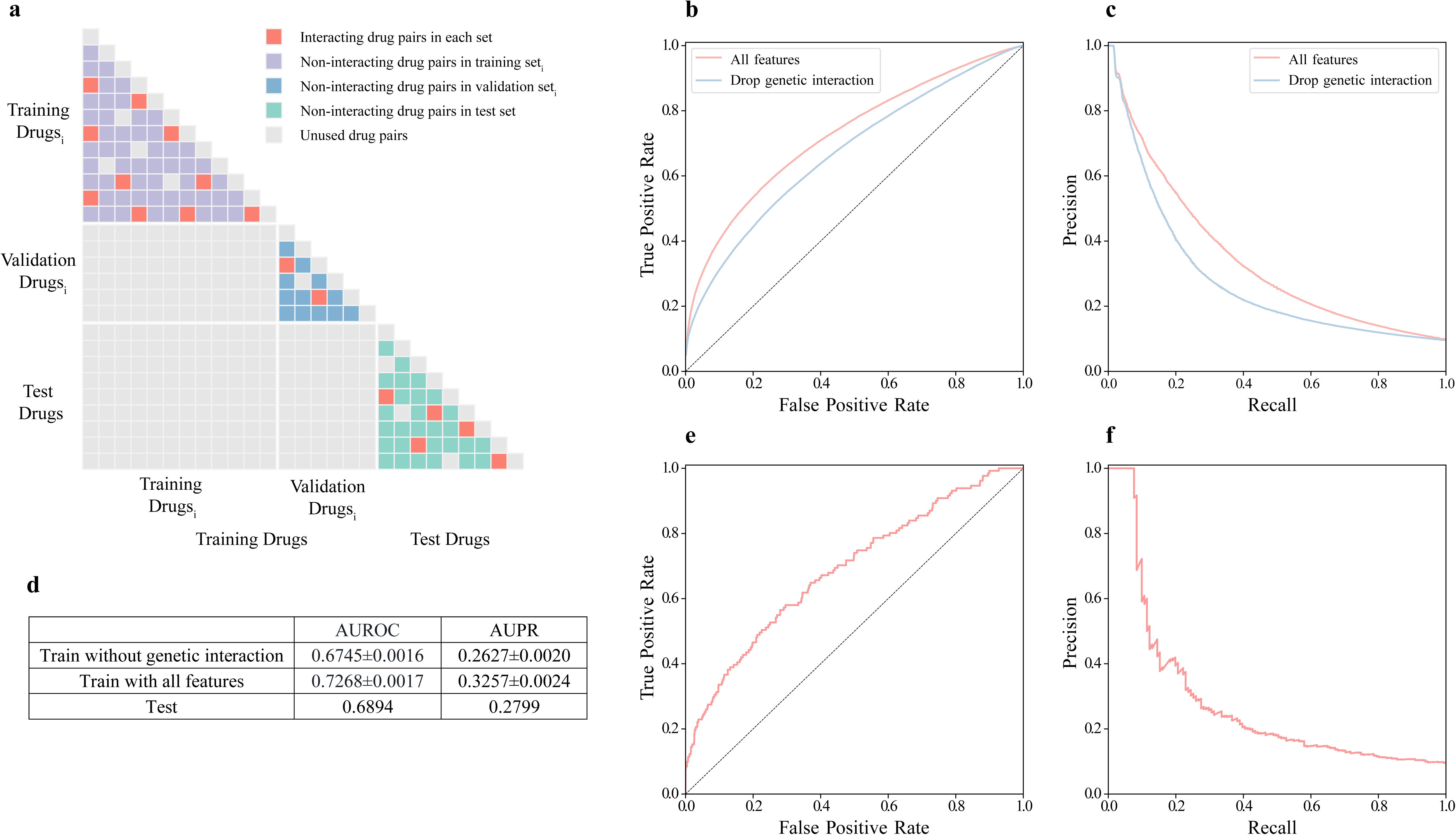
The train-test splitting scheme and model performance on the test set. (a) The train-test splitting scheme. Drugs are randomly divided into “training drugs” and “test drugs” with ratio of 2:1. Training set only consists of drug pairs constituted by “training drugs” and test set only consists of drug pairs constituted by “test drugs”. Training drugs are further split into “training drugs_i_” and “validation drugs_i_” with the same splitting scheme to obtain training set_i_ and validation set_i_ in the training phase. For each iteration of hold-out validation, the classifier is fit with training set_i_ and evaluated with validation set_i_. Purple squares represent non-interacting drug pairs in training set_i_. Blue squares represent non-interacting drug pairs in validation set_i_. Green squares represent non-interacting drug pairs in test set. Red squares represent interacting drug pairs in each set. Grey squares represent unused drug pairs. (b) Approximate receiver operating characteristic (ROC) curves on the training set. (c) Approximate precision-recall curves on the training set. (d)AUROCs and AUPRs on the training set and the test set. (e) Receiver operating characteristic (ROC) curve on the test set. (f) Precision-recall curve on the test set.

To build a more interpretable model and speed up the training process, we applied a feature selection method known as group minimax concave penalty (MCP) ^38^ that have been previously employed on biological datasets ^39^. This resulted in a final group of 11 features whose values were all significantly different between adversely interacting drugs and non-interacting drugs **(Fig. 1b-c, e-f**). An extreme gradient boosting (XGBoost) classifier ^40^ was then built because of its speed and outstanding performance in data science competitions. We optimized hyperparameters of the classifier using the tree-structured Parzen Estimator (TPE) approach ^41^, which has been shown to drastically improve the performance in a recent study predicting protein-protein interaction interfaces ^42^. Notably, instead of doing cross-validation, we adopted the same drug-based splitting scheme on the training set for hold-out validation (**Fig. 2a**). This enables the model to be best tuned for predicting interacting drug pairs without any prior information about the interaction profiles of the drugs involved. Indeed, a previous report by Liu et al. showed that classifier performance dropped significantly when evaluated on a test set consisted of pairs of drugs completely unseen in the training set if conventional cross-validation was performed ^21^. Our novel training strategy resulted in an average area under the receiver operating characteristic curve (AUROC) of 0.727 and an average area under the precision-recall curve (AUPR) of 0.326 over 1,000 trials of hold-out validation on the training set (**Fig. 2b-d**). When evaluated on the test set, our classifier achieved an AUROC of 0.689 (**Fig. 2d-e**) and an AUPR of 0.280 (**Fig. 2d, 2f**), demonstrating the robustness of our model. As shown in Table 1, our classifier attained a precision of 100% on the top 10 predictions, and a precision of 65% on the top 20 predictions (**Table 1**). Since there is no gold-standard set of non-interacting drugs, it is plausible that our non-interacting drug pairs might actually contain adverse DDIs. Not surprisingly, some non-interacting drug pairs with the high predicted probabilities can be found with evidence supporting their possible adverse interactions. For example, the drug pair with a non-interacting label with the highest predicted interacting probability in the test set, liothyronine and tretinoin, has been indicated to potentially cause intracranial pressure increase and a higher risk of pseudotumor cerebri when taken together^43^. Furthermore, diazoxide and spironolactone, predicted with an interacting probability of 0.846, have been reported to induce asthma, cardice hypertrophy and pulmonary edema according to FDA reports when co-administrated ^44^.

**Table 1.** Top 20 DDI predictions in the test set.

| ID1 | ID2 | Drug name1 | Drug name2 | Label | Probability |
| --- | --- | --- | --- | --- | --- |
| DB01076 | DB01098 | Atorvastatin | Rosuvastatin | 1 | 0.9943 |
| DB01076 | DB01095 | Atorvastatin | Fluvastatin | 1 | 0.9938 |
| DB00381 | DB00421 | Amlodipine | Spirolactone | 1 | 0.9889 |
| DB00880 | DB00887 | Chlorothiazide | Bumetanide | 1 | 0.9704 |
| DB00313 | DB01356 | Valproic Acid | Lithium cation | 1 | 0.9614 |
| DB00887 | DB00999 | Bumetanide | Hydrochlorothiazide | 1 | 0.9463 |
| DB00880 | DB01119 | Chlorothiazide | Diazoxide | 1 | 0.8835 |
| DB00421 | DB01076 | Spirolactone | Atorvastatin | 1 | 0.8815 |
| DB00421 | DB00622 | Spirolactone | Nicardipine | 1 | 0.8753 |
| DB00999 | DB01119 | Hydrochlorothiazide | Diazoxide | 1 | 0.8723 |
| DB00279 | DB00755 | Liothyronine | Tretinoin | 0 | 0.8677 |
| DB00162 | DB00755 | Vitamin A | Tretinoin | 1 | 0.8610 |
| DB00477 | DB01098 | Chlorpromazine | Rosuvastatin | 0 | 0.8601 |
| DB00421 | DB01119 | Spirolactone | Diazoxide | 0 | 0.8457 |
| DB00313 | DB00523 | Valproic Acid | Alitretinoin | 0 | 0.8419 |
| DB05015 | DB06176 | Belinostat | Romidepsin | 0 | 0.8378 |
| DB01065 | DB01069 | Melatonin | Promethazine | 1 | 0.8342 |
| DB00622 | DB01076 | Nicardipine | Atorvastatin | 1 | 0.7995 |
| DB00755 | DB00900 | Tretinoin | Didanosine | 0 | 0.7962 |
| DB00162 | DB01212 | Vitamin A | Ceftriaxone | 0 | 0.7858 |

To demonstrate the utility of our method, we obtained 5,039 drug pairs involving 295 drugs that had not been used for training and testing (**Supplementary Fig. 6**). After refitting our model on all 12,426 drug pairs that were used to develop our method, we predicted 432 novel DDIs (**Supplementary Table 2**). Remarkably, out of the top 20 newly predicted adversely interacting drug pairs, 9 can be verified in the TWOSIDES database (**Table 2**), manifesting the reliability of our method.

**Table 2.** Top 20 new adverse DDI predictions.

| ID1 | ID2 | Drug name1 | Drug name2 | Probability | In Twosides |
| --- | --- | --- | --- | --- | --- |
| DB00347 | DB01189 | Trimethadione | Desflurane | 0.9750 | No |
| DB00136 | DB00630 | Calcitriol | Alendronic acid | 0.9739 | Yes |
| DB00417 | DB01050 | Phenoxymethylpenicillin | Ibuprofen | 0.9699 | Yes |
| DB00228 | DB00347 | Enflurane | Trimethadione | 0.9680 | No |
| DB00347 | DB00753 | Trimethadione | Isoflurane | 0.9668 | No |
| DB00347 | DB01236 | Trimethadione | Sevoflurane | 0.9656 | No |
| DB00887 | DB01586 | Bumetanide | Ursodeoxycholic acid | 0.9567 | Yes |
| DB00532 | DB01189 | Mephenytoin | Desflurane | 0.9531 | No |
| DB00532 | DB01236 | Mephenytoin | Sevoflurane | 0.9521 | No |
| DB00228 | DB00532 | Enflurane | Mephenytoin | 0.9466 | No |
| DB01067 | DB01083 | Glipizide | Orlistat | 0.9456 | Yes |
| DB00731 | DB01016 | Nateglinide | Glyburide | 0.9430 | Yes |
| DB01050 | DB01053 | Ibuprofen | Benzylpenicillin | 0.9422 | No |
| DB00532 | DB00753 | Mephenytoin | Isoflurane | 0.9368 | No |
| DB00421 | DB01216 | Spironolactone | Finasteride | 0.9333 | Yes |
| DB00162 | DB00165 | Vitamin A | Pyridoxine | 0.9260 | No |
| DB01236 | DB04930 | Sevoflurane | Permethrin | 0.9250 | No |
| DB00284 | DB00731 | Acarbose | Nateglinide | 0.9199 | Yes |
| DB00136 | DB00273 | Calcitriol | Topiramate | 0.9132 | Yes |
| DB00877 | DB01586 | Sirolimus | Ursodeoxycholic acid | 0.9122 | Yes |

### Genetic interaction provides mechanistic insight into drug-drug interactions

We investigated the contribution of genetic interaction features to classifier performance by building and tuning a new model without them. Excluding genetic interaction features significantly decreases classifier performance when either AUROC or AUPR is examined (**Fig. 2b-d**). This establishes genetic interaction as an important feature in our model for predicting DDIs, providing complementary information that other features cannot capture.

More importantly, genetic interaction can help us generate plausible mechanistic explanations for drug-drug interactions. For example, mesalazine and dexamethasone, both of which are anti-inflammatory drugs, are a pair of drugs in the test set that have been labeled as adversely interacting. Mesalazine can target the IKBKB protein, whereas dexamethasone can target NOS2, which plays important roles in nitric oxide signaling. In yeast, double knockout of *ATG1* and *TAH18*, the respective yeast homologs of *IKBKB* and *NOS2*, exhibits a more negative impact on cell viability than expected from single knockout phenotypes ^35^. In human, IKBKB can phosphorylate the NF-κB inhibitor and activate NF-κB ^45^, which is a family of transcription factors involved in inflammation and immunity. Notably, the transcription of *NOS2* is induced by NF-κB activity ^46^. Mesalazine has been shown to inhibit IKBKB, thereby inhibiting the activation of NF-κB, while dexamethasone is a negative modulator of NOS2. A previous study has reported that dexamethasone can decrease NOS2 translation and facilitate NOS2 degradation in rat ^47^ (**Fig. 3a**). The combined use of mesalazine and dexamethasone may largely reduce the amount of NOS2, potentially affecting neurotransmission, antimicrobial and antitumoral activities.

**Figure 3.**
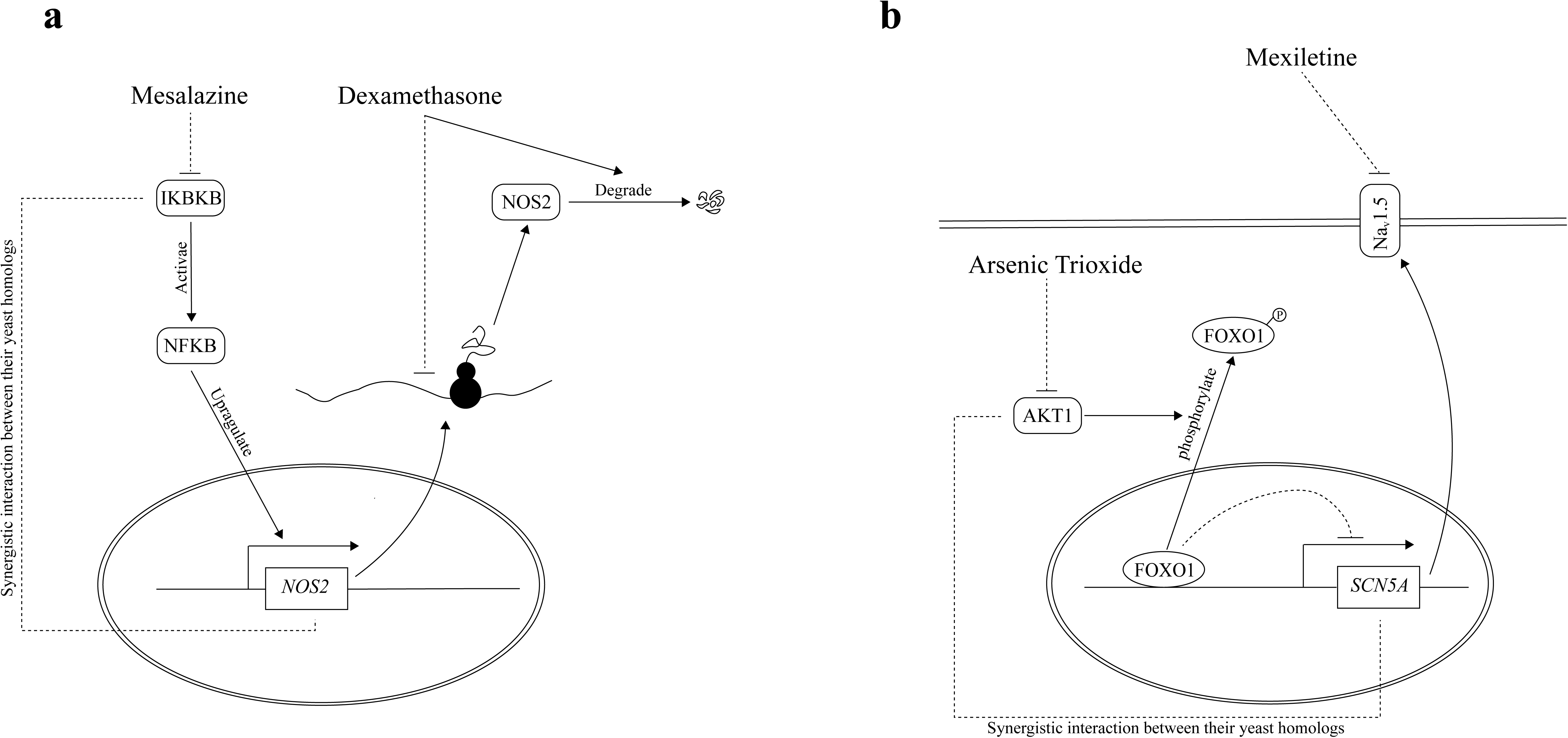
Genetic interaction provides possible mechanistic insights into DDIs. (a) Mesalazine inhibits IKBKB, a positive regulator of NF-κB activity, and NF-κB is a transcription factor which induces *NOS2* transcription. Dexamethasone can inhibit the transcription of *NOS2* and facilitate degradation of NOS2. The combined use of dexamethasone and mesalazine could potentially reduce the amount of NOS2 in cells to a large extent, which may affect neurotransmission, antimicrobial and antitumoral activities. (b) Mexiletine targets NAv1.5, a sodium channel encoded by *SCN5A*, while arsenic trioxide targets AKT1. The transcription of *SCN5A* is repressed by the transcriptional repressor FOXO1. AKT1 can activate the transcription of *SCN5A* by phosphorylating FOXO1. The combined use of mexiletine and arsenic trioxide could inactivate the transcription of *SCN5A* and at the same time block the existing sodium channel, which may largely reduce sodium influx in cardiac cells.

As another example, arsenic trioxide and mexiletine are a pair of drugs not labelled as adversely interacting in DrugBank, but predicted by our model to interact with high probability. As a chemotherapy drug for acute promyelocytic leukemia (APL), arsenic trioxide has been reported to decrease the activity of a serine/threonine-protein kinase AKT1 ^48^. On the other side, mexiletine is a sodium channel blocker that has also been used as part of a prophylactic therapy to treat APL patients to reduce cardiac complications ^49^. *PKC1*, the yeast homolog of *AKT1*, exhibits strong synergistic interaction with *CCH1* ^35,50^, which is the homolog of *SCN5A*, the gene encoding the sodium channel NA_v_1.5 targeted by mexiletine. In human, the transcription of *SCN5A* is repressed by FOXO1, whose transcriptional repression activity is in turn inactivated by AKT1-dependent phosphorylation ^51^ (**Fig. 3b**). Therefore, the simultaneous inhibition of AKT1 and the sodium channel by the two drugs may reduce sodium influx in cardiac cells to a greater extent, potentially causing undesired adverse effects. Indeed, this pair of drugs is reported by TWOSIDES as interacting, providing additional supporting evidence to their adverse interaction.

## DISCUSSION

In the past decade, many methods have been developed for predicting DDIs based on various types of features. In this study, we have incorporated a novel feature, namely genetic interaction, to build a gradient boosting-based model for fast and accurate adverse DDI prediction. We have shown that our classifier can robustly predict drug-drug interactions even for drugs whose interaction profiles are completely unseen during training. Furthermore, we have predicted 432 novel DDIs, with additional evidence supporting our top predictions, demonstrating the usefulness of our approach.

Most previous efforts of predicting DDIs suffer from an inability to make predictions for newly developed drugs due to train-test split based on drug pairs rather than drugs ^10,16,17,19,22,23,36,37^. Three studies attempted to address this problem by dividing the entire dataset based on drugs ^14,20,21^. However, they failed to do so during the training phase, resulting in an inflated performance on the training set. We have followed the drug-based train-test splitting scheme and have adopted a hold-out validation approach to avoid using overlapping drug sets for fitting the model and evaluating its performance. By doing so, we have achieved robust performance on the training set and the test set, which establishes the ability of our method to predict new DDIs for drugs whose interaction profiles are completely unknown.

By examining genetic interactions, our method provides mechanistic insights into how two drugs may interact in a detrimental fashion. The combined modulatory effect resulted from binding of two drugs to their respective targets might underlie adverse DDIs, and genetic interaction gives valuable information about the nature of such combined effect. Indeed, we have observed that genetic interaction features are indispensable to our classifier performance. Nevertheless, our work is limited by the lack of a global human genetic interaction network. As a surrogate for human genetic interactions, genetic interactions of yeast homologs were used in this study. Fortunately, large-scale human genetic interaction studies are coming into sight. Using a recently published dataset of human genetic interactions encompassing 222,784 gene pairs ^52^, we have found that the distribution of human genetic interaction scores vary significantly between adversely interacting drugs and non-interacting drugs (**Supplementary Fig. 2b**). With the continuous advancement of technologies for probing human genetic interactions including CRISPR interference, we anticipate that a more comprehensive map of human genetic interactions will become available in the near future, which could illuminate adverse DDI prediction to a larger extent.

## Acknowledgments

The authors would like to thank G. Hooker, S. Chen and S. Wierbowski for helpful discussions. This work was supported by National Institute of General Medical Sciences grants (R01 GM124559, R01 GM125639) and National Science Foundation grant (DBI-1661380) to H.Y.

## Author Contributions

H.Y. conceived the study and oversaw all aspects of the study. S.Q. and S.L. performed computational analyses and wrote the manuscript with input from H.Y. S.Q. generated figures and tables with input from S.L. and H.Y. All authors edited and approved of the final manuscript.

---

**Supplementary Figure 1.** Schematics of our DDI prediction framework. Four groups of features were calculated for each drug pair. Drug pairs were then divided into a training set and a test set. A gradient boosting-based model was built on the training set after feature selection. Model performance was evaluated on training set using hold-out validation and also on the test set. We demonstrate the importance of our novel feature with a case study and provide novel DDI predictions at the end.

**Supplementary Figure 2.** The distribution of adversely interacting drug pairs and non-interacting drug pairs in terms of the 5 unused features. (a) Indication similarity score of hierarchy level HLT between two drugs. (b) Side effect similarity score of hierarchy level PT and SOC between two drugs. (c) Mean and median genetic interaction score between targets of two drugs. (Statistical significance determined by two-sided Mann-Whitney U test)

**Supplementary Figure 3.** (a) The total number of protein targets between two drugs. (b) Minimum, mean, median and maximum human K562 cell line genetic interaction score between targets of two drugs. (Statistical significance determined by two-sided Mann-Whitney U test)

**Supplementary Figure 4.** The correlation between genetic interaction features and other features.

**Supplementary Figure 5.** Values of hyperparameters of the XGBoost model over 2000 TPE iterations.

**Supplementary Figure 6.** Construction of a set of drug pairs used for new predictions. All combinations between drugs that appear in the first category in DrugBank and other drugs, as well as all pairwise combinations of drugs not in the first category, are taken for new predictions. Green squares represent drug pairs used for building the classifier. Grey squares represent unused drug pairs. Blue squares represent drug pairs used for new predictions.

**Supplementary Table 1.** Five main DDI categories in DrugBank.

**Supplementary Table 2.** A list of 432 new adverse DDI predictions.

## ONLINE METHODS

### Data collection

We obtained DDI data from DrugBank (version 5.0.10) ^28^. Among the 5 major interaction categories in DrugBank (**Supplementary Table 1**), we only considered the first category as they were clearly defined as adverse DDIs. Non-interacting drug pairs were constructed by taking all other combinations using the same set of drugs, removing drug pairs also appearing in other categories in DrugBank, TWOSIDES ^29^, or a complete dataset of DDIs ^30^ compiled from a number of sources. This minimizes the chance of having actual adverse DDIs in the non-interacting set given the absence of a gold standard set of non-interacting drug pairs. From DrugBank, we also collected human protein targets of drugs and their sequences.

Side effects were obtained from SIDER 4.1 ^53^ and OFFSIDES ^29^. Both databases use UMLS concept IDs as their side effect identifiers. However, as reported by Zhang et al. ^20^, some side effect terms are similar, and synonyms could cause biases when calculating side effect similarity. To solve this problem, we obtained mapping from UMLS concept IDs to MedDRA concept IDs from the 2017AB release of UMLS ^54^. Furthermore, we obtained the full MedDRA hierarchy from MedDRA (version 21.0) ^31^. This allowed us to map UMLS concept IDs to different levels (PT, HLT, HLGT and SOC) of the MedDRA hierarchy. Similar to side effect data, indications of drugs were acquired from SIDER 4.1 ^53^ and mapped to the same 4 levels of the MedDRA hierarchy.

For genetic interactions, we obtained yeast genetic interactions from Costanzo et al. ^35^. We first filtered all genetic interactions by a p-value cutoff of 0.05 and aggregated the scores of all combinations of alleles of each yeast gene pair by applying the arithmetic mean. Drug targets in the form of UniProt IDs were mapped to gene names by UniProt ^55^ and these human genes were mapped to their yeast homologs via SGD YeastMine ^56^. For human gene pairs mapped to multiple yeast gene pairs, we obtained a single score for each human gene pair by applying the arithmetic mean.

### Feature extraction and the train-test split

For a drug pair (A,B), four groups of features were calculated (**Fig. 1a, d**): indication similarity scores between A and B, side effect similarity scores between A and B, target sequence similarity scores between targets of drug A and targets of drug B, and genetic interaction scores between targets of drug A and targets of drug B. Indications and side effects of drugs were mapped to 4 different levels of the MedDRA hierarchy as described above. At each level, indication similarity was calculated by taking the Jaccard index between the respective indication vectors of drug A and drug B (**Fig. 1a**). Similarly, side effect similarity was calculated by applying the same measure on the side effect vectors at the 4 different MedDRA hierarchy levels (**Fig. 1a**). For genetic interactions, since each drug can have multiple targets, we obtained a single score for each drug pair by aggregating the genetic interaction scores of all their corresponding target pairs using 4 different functions, namely taking the minimum, mean, median or maximum (**Fig. 1d**). Similarly, the same 4 functions were used for constructing target similarity features, which was calculated from the target sequences with the Smith-Waterman algorithm using the scikit-bio Python library. The raw scores were normalized as described in Bleakley et al. ^57^. Overall, 16 features belonging to 4 feature groups were constructed. Only drug pairs with all features available were considered when building the machine learning model. All drugs were randomly split into “training drugs” and “test drugs” with a 2:1 ratio. The training set consisted of all drug pairs where both drugs were “training drugs” and the test set consisted of all drug pairs where both drugs were “test drugs” (**Fig. 2a**). We constrained the fraction of adversely interacting drug pairs in the training set and that in the test set to be fairly balanced. To obtain the optimal feature combination, we calculated all features for the training set and applied group minimax concave penalty (MCP) ^38^ with the ‘grpreg’ R package with default parameters. All subsequent training was done using this optimal set of features.

### Hyperparameter optimization and classifier training

The gradient boosting-based algorithm XGBoost ^40^ was used in this study. To find the best combination of hyperparameters for the XGBoost classifier, the tree-structured Parzen estimator (TPE) approach ^41^ was adopted. Because of the drug-based approach by which we split our dataset into training and test sets, we applied the same splitting scheme on the training set multiple times to obtain training set_i_ and validation set_i_ instead of simply using cross-validation. Each split on the training set can be seen as a hold-out validation, as we used training set_i_ to fit the model and validated model performance on validation set_i_. We selected the average AUPR of 50 trials of hold-out validation as the loss function to minimize for TPE, and we ran TPE for 2,000 iterations to obtain set of hyperparameters that minimized the loss function for our XGBoost classifier (**Supplementary Fig. 5**). After finding the optimal set of hyperparameters, we fit the model on the complete training data.

### Model evaluation

Model performance on training set was evaluated by 1,000 runs of hold-out validation on the training set. For each hold-out validation, we fitted the model on training set_i_ and obtained AUROC and AUPR. We averaged AUROC and AUPR over 1,000 runs of hold-out validation as measurements of the performance of the model. Approximate ROC curve and precision-recall curve (**Fig. 2b-c**) were plotted by averaging the 1,000 ROC curves and 1,000 precision-recall curves respectively at every thousandth of a point on the x-axis. In order to evaluate the ability of the classifier to identify drug-drug interactions between drugs whose interaction profiles were completely unknown during training, the model was evaluated on the test set which had no overlap with the training set in terms of the drugs involved. Predictions were ranked according to their raw prediction scores to produce the ROC curve and the precision-recall curve.

### Making new predictions

To make novel adverse DDI predictions, we examined all combinations of drugs that appeared in DrugBank, excluding drug pairs where both drugs were involved in the first category of DDIs (**Supplementary Fig. 6**), which we used for building the machine learning model. We then predicted 6,690 drug pairs involving 336 drugs for which all features could be calculated using the classifier retrained on the whole dataset. The probability cutoff that produced the maximum averaged F1 score over 1,000 runs of hold-out validation on the training set was chosen for determining new DDI predictions.

