## Supplementary Figures and Supplementary Table 1 for "Leveraging genetic interaction for adverse drug-drug interaction prediction"

### Supplementary Figure 1

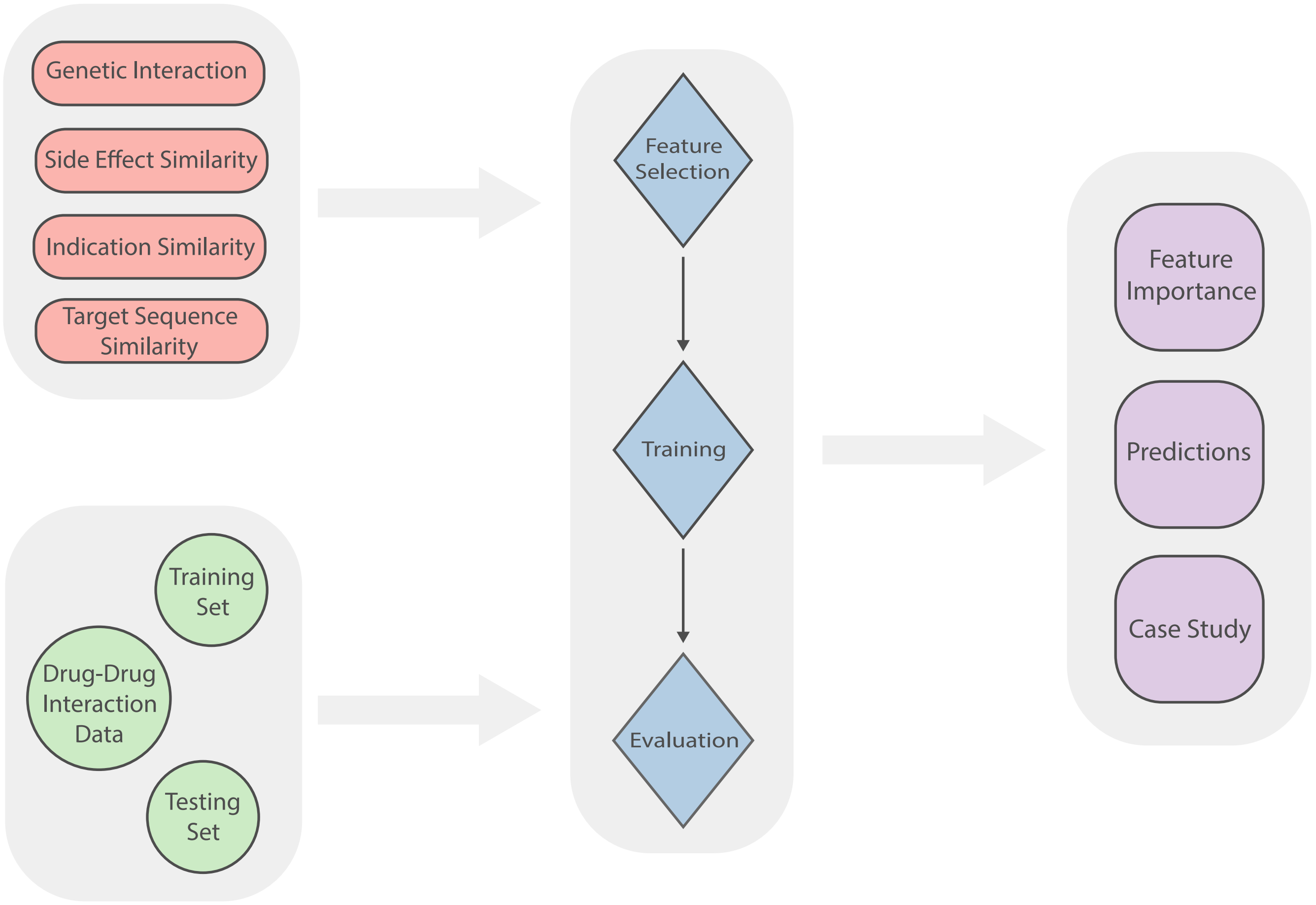

### Supplementary Figure 2

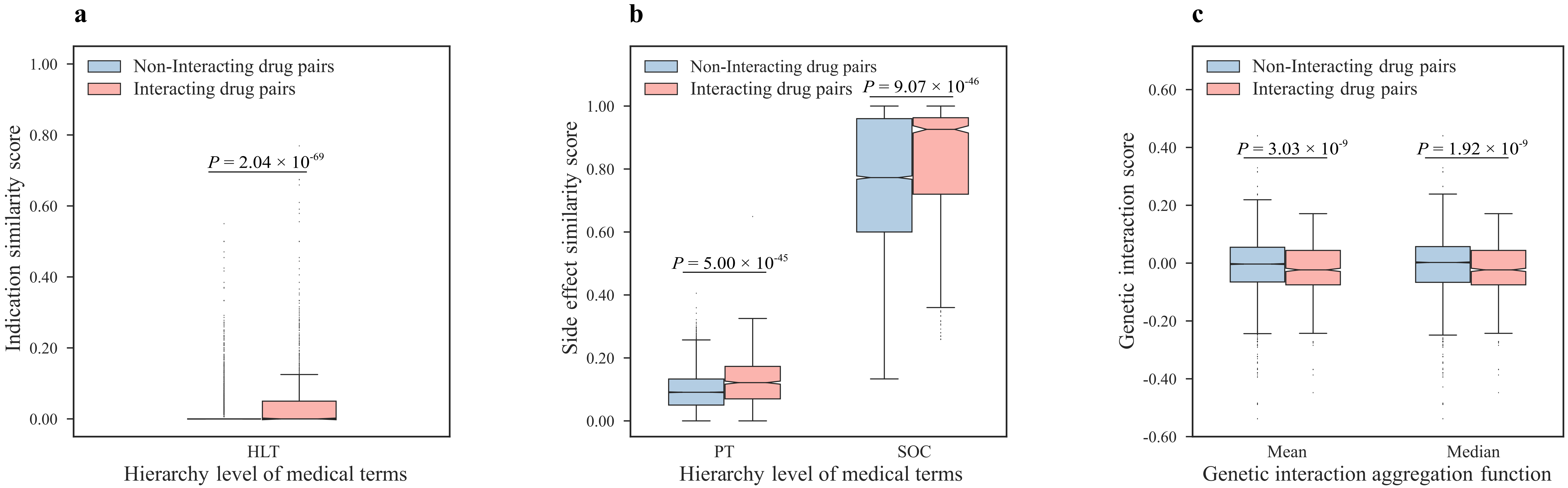

#### Supplementary Figure 3

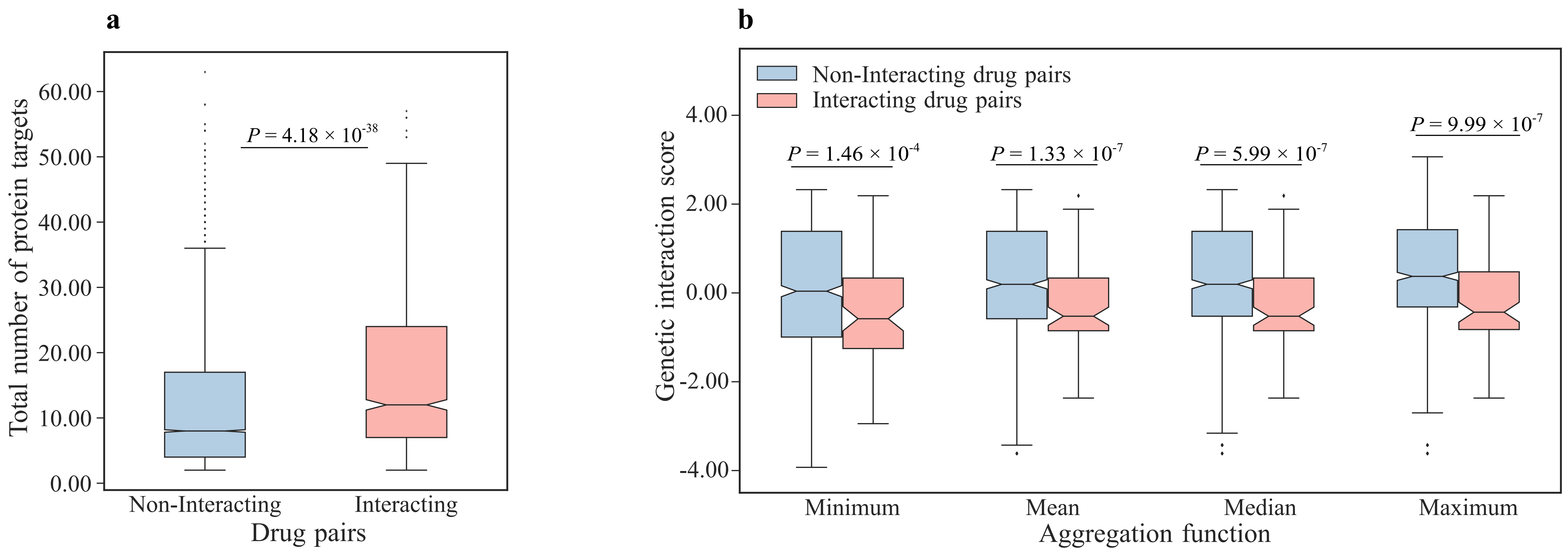

### Supplementary Figure 4

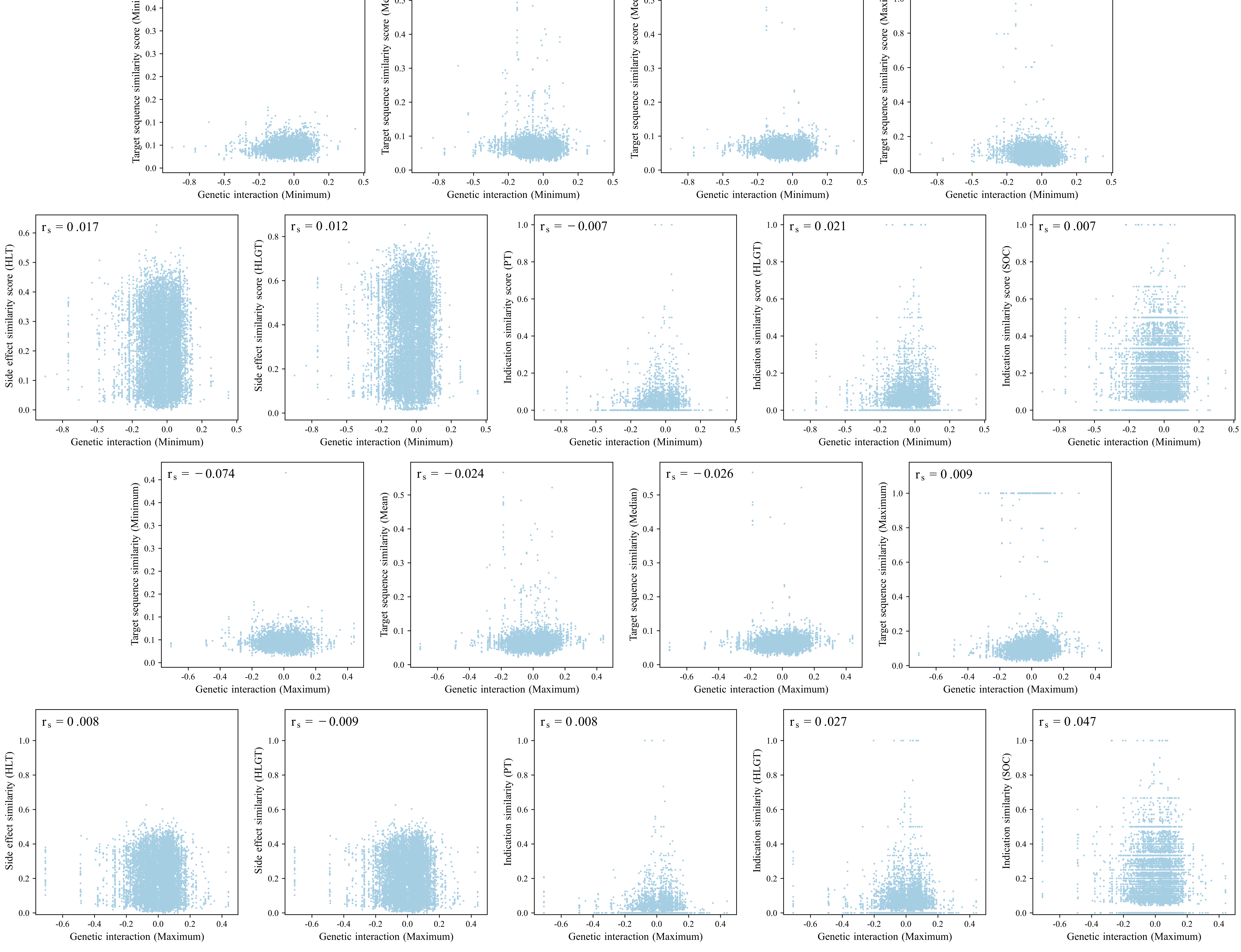

### Supplementary Figure 5

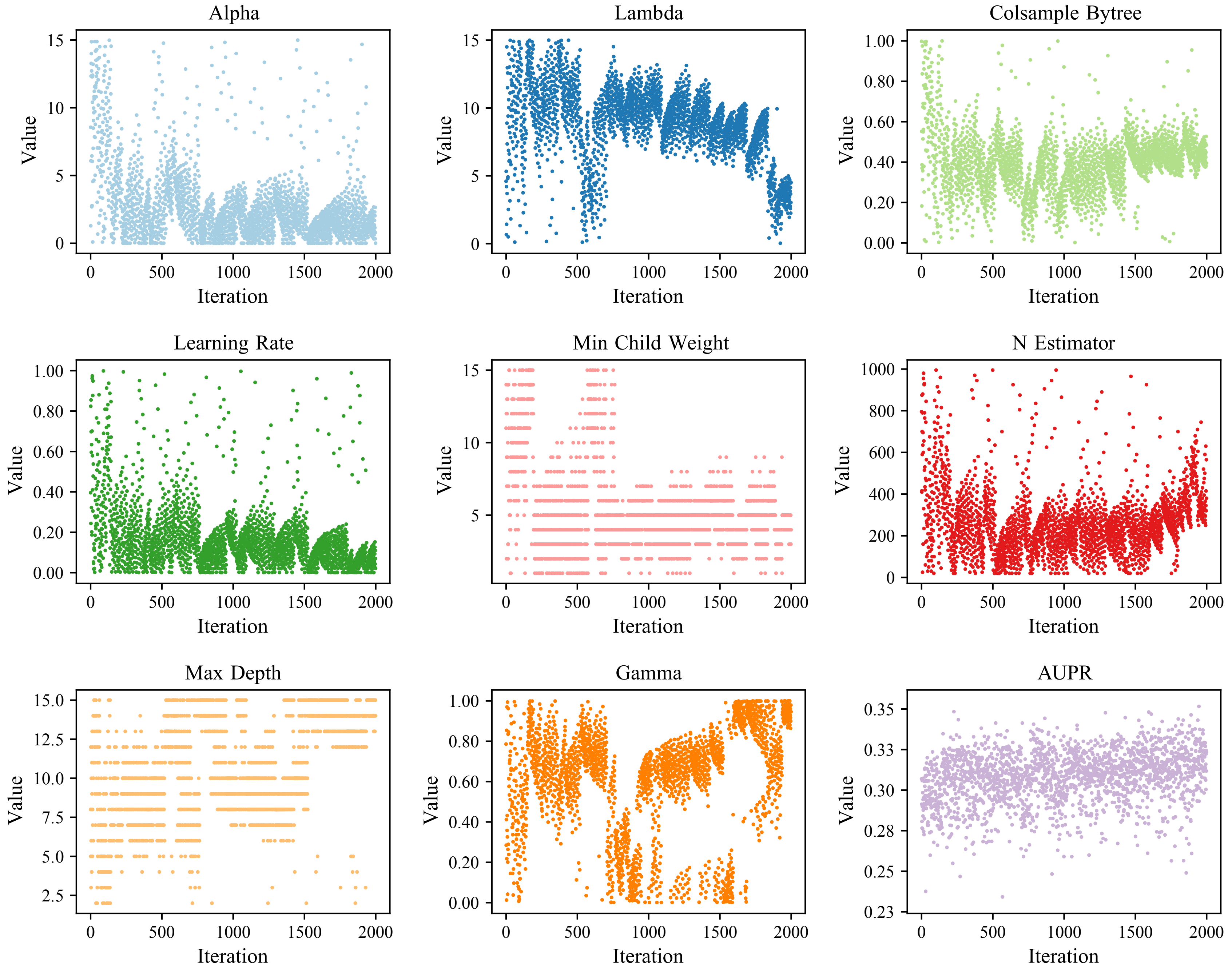

### Supplementary Figure 6

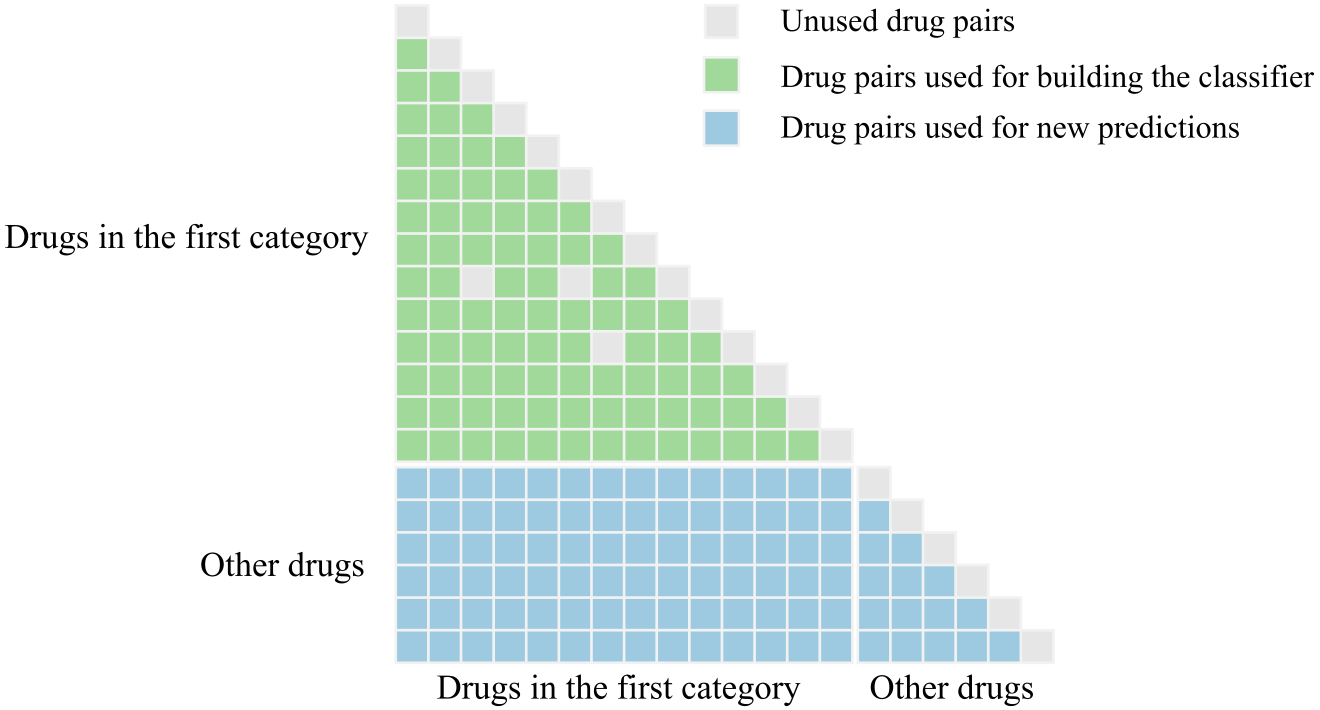

### Supplementary Table 1

| Category | Amount |
|---|---|
| The risk or severity of adverse effects can be increased when Drug A is combined with Drug B. | 117045 |
| Drug A may decrease/increase the (...) activities of Drug B. | 100440 |
| The serum concentration of Drug A can be increased/decreased when it is combined with Drug B. | 73478 |
| The metabolism of Drug A can be increased/decreased when combined with Drug B. | 41999 |
| The therapeutic efficacy of Drug A can be decreased when used in combination with Drug B. | 22358 |
